# Dynamic neural states underpin bradykinesia severity in Parkinson’s disease

**DOI:** 10.1101/2025.08.26.672349

**Authors:** Abhinav Sharma, Tao Liu, Bahman Abdi-Sargezeh, Amelia Hahn, Maria Scherbakova, Wolf-Julian Neumann, Simon Little, Philip Starr, Ashwini Oswal

## Abstract

**Background:** Bradykinesia in Parkinson’s disease (PD) may arise due to transient, network-wide neural dynamics that extend beyond beta-band oscillatory activity within the motor cortical-subthalamic nucleus (STN) circuit.

**Methods:** We address this question by using Hidden Markov Models (HMMs) to identify neural states from chronic motor cortical and STN recordings in five PD patients (1,046 hours from 10 hemispheres), with concurrent measurements of bradykinesia using wearable sensors.

**Findings:** We identified four neural states with distinct spectral and temporal features. Two states exhibited spectral signatures—particularly STN low and high gamma, STN delta/alpha, cortical beta, and cortico-STN beta coherence—that predicted worsening bradykinesia. However, STN beta power alone was not consistently predictive, challenging traditional beta-centric views. These states also displayed compensatory features associated with bradykinesia amelioration, including cortical delta/alpha activity, cortical high gamma, and cortico-STN high gamma coherence. Two additional states affected bradykinesia through temporal rather than spectral properties. Prolonged lifetimes of one of these state worsened symptoms, whereas increased occurrences of another, marked by local beta without cortico-STN beta coherence, improved motor function.

**Interpretation:** Our findings highlight the multidimensional nature of bradykinesia and suggest that state-aware, adaptive interventions targeting state features—rather than single frequency bands—may offer new opportunities for improved deep brain stimulation in PD.

**Funding:** AO is supported by an MRC Clinician Scientist Fellowship (MR/W024810/1) and a Rosetrees Trust/Race Against Dementia Team award. BA and AO acknowledge funding support from the Oxford University Hospitals Charity and Jon Moulton Trust. TL acknowledges funding support from the China Scholarship Council.

**Research in Context:** *Evidence before this study:* Previous work has demonstrated that subthalamic nucleus oscillatory activity at beta (15-30 Hz) frequencies correlates positively with motor symptoms in Parkinson’s disease. This has led to beta activity being used as a biomarker for adaptive Deep Brain Stimulation. It remains unclear however whether other oscillatory features within the broader motor cortical-subthalamic nucleus network could provide improved biomarkers for tracking symptom severity.

*Added value of this study:* We address this by performing chronic motor cortical and subthalamic nucleus recordings in Parkinson’s disease patients during activities of daily living. Simultaneous measurements of symptom severity were captured using wearable sensors. We used Hidden Markov Models to identify transient states of neural activity and related these to symptom severity. Although cortical beta and cortico-STN beta coherence predicted worsening motor symptoms, STN beta activity alone was not a consistent predictor. Interestingly, we identified spectral features associated with motor symptom improvements, including cortical delta/alpha activity, cortical high gamma, and cortico-STN high gamma coherence. Additionally, there was a ‘compensatory’ state characterised by short-lived cortical and subthalamic nucleus beta activity, whose increased occurrence was associated with symptomatic improvements.

*Implications of all the available evidence:* Our findings highlight the importance of prolonged, high temporal resolution measurements of both neural activity and symptom severity for discovering adaptive Deep Brain Stimulation biomarkers. Crucially, we identify new target states and spectral features for improving motor symptoms in Parkinson’s disease.

## Introduction

Bradykinesia is a cardinal motor symptom of Parkinson’s disease (PD), characterized by a pathological reduction in movement velocity and amplitude, significantly impairing patients’ quality of life ^1–3^. From a therapeutic perspective, bradykinesia exhibits a robust response to dopaminergic therapy and high-frequency deep brain stimulation (DBS) ^4–6^. Mechanistically, beta-band oscillatory activity (12–30 Hz) within the subthalamic nucleus (STN) has been extensively investigated as a neurophysiological correlate of bradykinesia, often assessed through the motor subsection of the Unified Parkinson’s Disease Rating Scale (UPDRS-III)^7–15^. At the network level, increased coherence between the motor cortex and STN within the beta frequency range has been implicated in akinetic-rigid manifestations of PD, suggesting a circuit-wide pathophysiological mechanism^16,17^. Notably, cortical beta activity, particularly within primary motor regions, has also been linked to worsening motor deficits in PD, reinforcing the hypothesis that exaggerated beta synchrony contributes to impaired motor function^18–22^. Both dopaminergic medication and DBS can attenuate beta oscillatory activity within the cortex and STN, while concurrently reducing cortico-subthalamic synchrony. These neurophysiological changes closely correlate with improvements in motor function, establishing a strong association between pathological beta synchronization and bradykinesia^10,23–27^.

Recent research has increasingly recognized the pathological significance of oscillatory dynamics beyond the beta band, implicating multiple frequency domains in the pathophysiology of PD^28,29^. Furthermore, some evidence also challenges the traditional view that beta oscillations alone drive bradykinesia, as experimental manipulations of beta-band activity in animal models do not consistently exacerbate motor impairment^30–32^. Recently it has been proposed that low-frequency rhythms in the delta (∼2–4 Hz) and theta (∼4–10 Hz) bands contribute to motor dysfunction. Exaggerated low-frequency oscillations in the STN have been observed in dopamine-depleted states^33^, with correlations to bradykinesia and akinesia^33–35^.

In the higher frequency domain, gamma oscillations (40–100 Hz) have been consistently identified as exerting a prokinetic influence in Parkinson’s disease (PD). Investigations into gamma-band activity typically involve spectral power analyses in key regions, notably the subthalamic nucleus (STN) and motor cortex, as well as assessments of cortico-subthalamic coherence. These measures are frequently correlated with UPDRS-III motor scores under dopaminergic medication conditions^36–38^. Task–specific studies reinforce the prokinetic role of gamma oscillations, demonstrating their positive association with movement velocity, vigor, grip force, and their augmentation in response to levodopa administration^39–42^. Mechanistically, gamma activity is thought to function in direct opposition to beta oscillations in PD. This dynamic interplay suggests that an optimized motor state, characterized by diminished bradykinesia and rigidity, emerges when beta-band synchrony is suppressed, concomitant with an upregulation of gamma oscillations within the STN or the STN-motor cortex network^39,41^.

The motivation for this study stems from the growing realization that bradykinesia does not arise from a single, uniform neural substrate, but rather from dynamically fluctuating and spatially distributed circuit states involving diverse oscillatory regimes. While previous studies have established broad associations between frequency bands and motor symptoms, the underlying assumption of temporal stationarity and spatial homogeneity remains untested. Indeed, a central paradox in Parkinsonian physiology is that similar clinical manifestations may arise from distinct, and sometimes even opposing, spectral configurations - beta synchrony may co-occur with compensatory gamma bursts; delta oscillations may mark a persistent akinetic state or transiently decouple during gait. Static metrics such as averaged spectral power or global coherence obscure this underlying heterogeneity^43,44^.

To resolve this mechanistic opacity, we adopt a probabilistic framework grounded in Hidden Markov Models (HMMs) to characterize recurring spatiotemporal patterns,“states”, within continuous neural recordings^43^.These states capture the unique covariance structure of STN and motor cortical activity, as well as inter-regional coherence across canonical frequency bands. Crucially, HMMs allow for state identity to remain stable despite internal spectral variability, reflecting the biological plausibility of network configurations with multidimensional and partially dissociable components.

We apply HMMs to chronic neural activity recordings from DBS devices and correlate spectral and temporal features of the identified network states to dynamic measurements of bradykinesia from wearable sensors. Hereby, we leverage a framework that resolves moment-to-moment heterogeneity in local and circuit-level dynamics and reframes bradykinesia as a dynamic expression of latent pathological states, amenable to precise, state-targeted therapeutic interventions. This represents a shift away from one-dimensional biomarkers that dominate current clinical decision-making towards a multidimensional understanding of neural state architecture in PD.

## Methods

### Patient Cohort and Electrode Localization

We analysed data from five individuals diagnosed with Parkinson’s disease (PD), each implanted bilaterally with the Medtronic Summit RC+S neural interface. Subthalamic nucleus (STN) recordings were acquired via quadripolar Medtronic 3389 electrodes, while cortical signals were recorded using quadripolar paddle-type leads with 10 mm inter-contact spacing. The cortical leads were positioned to ensure that 2–3 contacts lay anterior to the central sulcus and approximately 2–4 cm lateral to the midline. Electrode localization was verified using a linear co-registration pipeline that fused postoperative CT with preoperative 3T MRI, following previously validated protocols^36,45^. Electrode contact coordinates were normalized to Montreal Neurological Institute (MNI) space using the Lead-DBS toolbox^46^ for group-level visualisation (**Supplementary Figure 1**).

Neural signals were streamed at a sampling rate of 250 Hz from both hemispheres to a Microsoft Windows tablet. Bipolar recordings were streamed from STN contacts 1–3 and 2–4 (numbers ordered from inferior to superior) and from cortical contacts 1–3 and 2–4 (numbers ordered from anterior to posterior). At least one STN contact was placed within the sensorimotor subregion of the STN, and one cortical contact overlayed the motor cortex (**Supplementary Figure 1**). Patients initiated home recordings were streamed to the Microsoft tablet in 1-2 week recording sprints (see Supplementary Methods and Gilron et al., 2021^45^).

Recordings were acquired 2–4 weeks post-surgery whilst patients performed naturalistic behaviours in their home environment, whilst taking their regular dopaminergic medication, prior to deep brain stimulation (DBS) initiation. Clinical characteristics of each patient are provided in **Table 1**. Concurrent motor symptom data were collected using bilateral wrist worn Personal KinetiGraph® (PKG®) monitors (Global Kinetics Pty Ltd.), which provided continuous two-minute interval assessments of bradykinesia, tremor, and dyskinesia, based on validated criteria^45^.

**Table 1.** Clinical characteristics of patients. LDE = levodopa dose equivalent. The total pre-operative UPDRS part III score is presented in the off-medication state.

| **Patient** | **Age and gender** | **Disease duration (years)** | **Preoperative medication (mg)** | **UPDRS III off medication** | **% Change in UPDRS III when on** | **MOCA** | **Data duration with concurrent contralateral PKG (hours)** |
| --- | --- | --- | --- | --- | --- | --- | --- |
| 1 02 | 54 M | 7 | LDE 1425 | 49 | 90% | 26 | L: 123.7<br>R: 192.9 |
| 2 05 | 63 M | 19 | LDE 955 | 45 | 51% | 30 | L: 27.7<br>R: 24.0 |
| 3 06 | 28 F | 12 | LDE 1550 | 61 | 73% | 27 | L: 80.5<br>R: 162.7 |
| 4 07 | 40 M | 4 | LDE 1314 | 41 | 65% | 30 | L: 192.7<br>R: 175.6 |
| 5 08 | 58 M | 12 | LDE 2100 | 44 | 75% | 27 | L: 40.4<br>R: 25.7 |

### Ethics

Ethical approval was granted by the University of California, San Francisco IRB. under a physician sponsored investigational device exemption (IDE), protocol # G180097, and was registered at ClinicalTrials.gov (NCT03582891). Signed informed consent was obtained from all participants.

### Hidden Markov Model Fit

A single Hidden Markov Model (HMM) was trained on the longitudinal neural recordings from all five patients. Each input file corresponded to one hemisphere and included four bipolar channels, two from the motor cortex and two from the STN. Files were excluded if any channel lacked valid signal or contained non-contiguous segments; in such cases, continuous fragments were isolated for analysis (see Methods: Patient Cohort and Electrode Localization). To reduce the influence of amplitude variability across channels, each channel was separately subjected to percentile normalization across the full dataset.

Time-delay embedding was applied using a symmetric 15-point window (−7:0:+7), transforming the 4-channel input into a 60-dimensional vector per time point (4 channels × 15 delays), spanning 60 ms at 250 Hz. Signals were bandpass filtered (2–120 Hz) to retain physiologically relevant oscillatory dynamics. The resulting data matrix (60 × time) for each session was used as input to the HMM, implemented using the Python version of the OSL-dynamics toolbox^47^ (https://osl-dynamics.readthedocs.io/en/latest/index.html). Each latent state was modelled using multivariate Gaussian emissions, initialized with random mean vectors and full-rank covariance matrices. The covariance matrices inferred from the estimation regime capture all relevant spectral features for each state. This covariance includes not just instantaneous correlations between regions (e.g., STN and cortex), but also cross-frequency phase relationships and time-lagged interactions that reflect both local rhythmicity and large-scale coupling patterns^43,48^. For instance, a given state might be defined by strong cortico-subthalamic coherence at beta frequency, STN-local autocovariance at a gamma timescale, or some hybrid of both. As long as these covariance features persist, the state identity remains intact, even if absolute spectral power fluctuates.

Crucially, this formulation enables a unified characterization of both local and circuit-level dynamics within a single probabilistic framework. Rather than treating power and coherence as separate variables or applying windowed averages, the TDE-HMM integrates them into a joint representation. This allows us to dissect how transient shifts in network-level synchrony or regional oscillatory bursts contribute to motor symptoms without collapsing them into a single frequency bin. Models with 3, 4, and 6 states were evaluated, and the 4-state model was selected for downstream analyses (see Spectral Characterization of HMM States for justification).

### PKG® Alignment and Preprocessing of Behavioral Data

PKG® bradykinesia scores were temporally aligned with neural recordings using timestamp synchronization^22,45^. Each PKG® score corresponded to a two-minute segment of neural data, defining the analysis window. For contralateral alignment, neural data from one hemisphere were paired with PKG® scores from the opposite wrist.

Bradykinesia scores were transformed to positive values for consistency. This meant that a higher bradykinesia score indicates worse bradykinesia in the patient. Epochs associated with prolonged immobility (duration >2 min and bradykinesia >80) were classified as sleep and excluded. To focus on data segments that captured a broad range of variance in bradykinesia scores, without including data that would be confounded by the presence of on state dyskinesias, we excluded epochs where bradykinesia scores were below each patient’s 30th percentile across the full recording period^36,45^.

### Static Spectral Analysis

For each two-minute window, we employed Welch’s method to calculate auto-power spectral densities (PSDs) for individual brain regions (STN and motor cortex) and cross-power spectral densities between regions. We applied a 50% overlap between analysis windows with a window length of 2 seconds and a Hann windowing function, yielding a frequency resolution of 0.5 Hz across the 2-100 Hz frequency range. This frequency resolution was consistent with our spectral analysis of HMM states. Magnitude squared coherence between STN, and motor cortex was calculated from the auto-and cross-PSDs.

### Spectral Characterization of HMM States

State time courses were extracted from the trained HMM, yielding continuous-valued probabilities (range: 0–1) representing the instantaneous likelihood of each state being active. These were subsequently binarized using the Viterbi algorithm to obtain discrete state assignments.

Within each two-minute window, state-specific neural data were isolated by element-wise multiplication of binarized state time courses with the corresponding non-embedded raw signals. Power spectral density (PSD) and coherence were computed using multi-taper spectral analysis with seven Slepian tapers (time-bandwidth product of 4, frequency resolution of 0.5 Hz) and 50% overlapping windows.

For each state, PSDs were averaged across STN channels to produce a single STN PSD, and across cortical channels to yield a CTX PSD. Coherence was computed for all STN–CTX channel pairs and averaged to derive a single STN–CTX coherence value per state. These features together formed the combined spectral and coherence fingerprint of each latent brain state.

In the standard OSL-dynamics pipeline, spectral outputs are often subjected to non-negative matrix factorization (NNMF) to reduce noise artifacts that arise due to limited sample sizes, a problem exacerbated by models with a large number of states or spatial channels. In our dataset, spatial dimensionality was not a limiting factor. However, training a six-state HMM degraded spectral estimation quality to the extent that NNMF became necessary. Because NNMF introduces additional complexity and requires parameter tuning, we opted for a four-state model. This reduction significantly improved the spectral clarity of states and obviated the need for NNMF, thereby simplifying the pipeline and enhancing interpretability for mechanistic clinical inference.

### Temporal Metrics of HMM States

Discrete state time courses allowed the calculation of several temporal features within each two-minute analysis window:

- Fractional Occupancy (FO): Proportion of time each state was active.
- State Lifetime: Average duration of continuous occupancy of a given state.
- Visit Interval: Mean interval between successive activations of the same state.

These metrics collectively characterized the temporal properties of latent brain states.

### Modelling Bradykinesia using Static Spectral Features

To establish the relationship between static spectral features and bradykinesia severity, we employed a general linear model (GLM) framework with two-minute time windows as units of analysis. For each window, we extracted 15 spatio-spectral features, comprising three spatial metrics (STN PSD, CTX PSD, and STN-CTX coherence) across five canonical frequency bands: delta-alpha (2-10 Hz), low-beta (12-20 Hz), high-beta (20-35 Hz), low-gamma (40-70 Hz), and high-gamma (70-100 Hz). This conventional approach computed spectral features by averaging across all neural configurations within each window.

The bradykinesia score at the conclusion of each window served as the dependent variable. Hemisphere and participant identity were modelled as categorical covariates to control for inter-hemispheric and inter-individual variability. P-values for individual predictors were corrected for multiple comparisons using FDR correction, with statistical significance evaluated at an alpha threshold of 0.01. Similar covariates and multiple comparisons correction were applied for all subsequent GLM analyses.

### Modelling Bradykinesia using HMM Derived Spatio-Spectral Features

As per the static spectral modelling, we leveraged a GLM, to interrogate the relationship between HMM derived spatio-spectral features and bradykinesia severity. For each 2-minute-long window, we extracted 15 spatio-spectral features per state, comprising three spatial metrics (STN PSD, CTX PSD, and STN–CTX coherence) across five canonical frequency bands: delta–alpha (2–10 Hz), low-beta (12–20 Hz), high-beta (20–35 Hz), low-gamma (40–70 Hz), and high-gamma (70–100 Hz). Across four states, this yielded 60 predictors.

### Modelling Bradykinesia using HMM Derived Temporal Features

Finally, GLM analyses were also deployed to probe the relationship between state temporal properties and bradykinesia severity. Given the inherent correlations between temporal metrics across states (e.g., fractional occupancies summing to unity), we fit separate GLMs for each temporal property from each state, to avoid multicollinearity. For each model, the dependent variable was the PKG® bradykinesia score, whilst the independent variable was the temporal feature of interest for each HMM state: fractional occupancy, mean lifetime, visit interval, or switching rate.

## Results

### Spectral Fingerprints of Cortico-Subthalamic States

We first sought to characterise the spatial and spectral properties of time resolved latent states inferred by the HMM and to compare these with time averaged (static) spectra. Each HMM state captured both local dynamics within individual brain regions (subthalamic nucleus and motor cortex) and the coherence patterns between these regions, enabling a comprehensive assessment of both local neural activity and inter-regional connectivity (**Figure 1**). **Figure 2** shows the results of spectral analysis for states 1-4, with STN power spectra (**Figure 2a**), motor cortical power spectra (**Figure 2b**) and STN-motor cortex coherence (**Figure 2c**) all displayed. The first column displays the static spectral characteristics of the entire dataset, enabling direct visual comparison between state-resolved and conventional spectral analysis. This comparison demonstrates that the state-resolved approach provides finer resolution about the spatial and temporal properties of cortico-STN network configurations.

**Figure 1.**
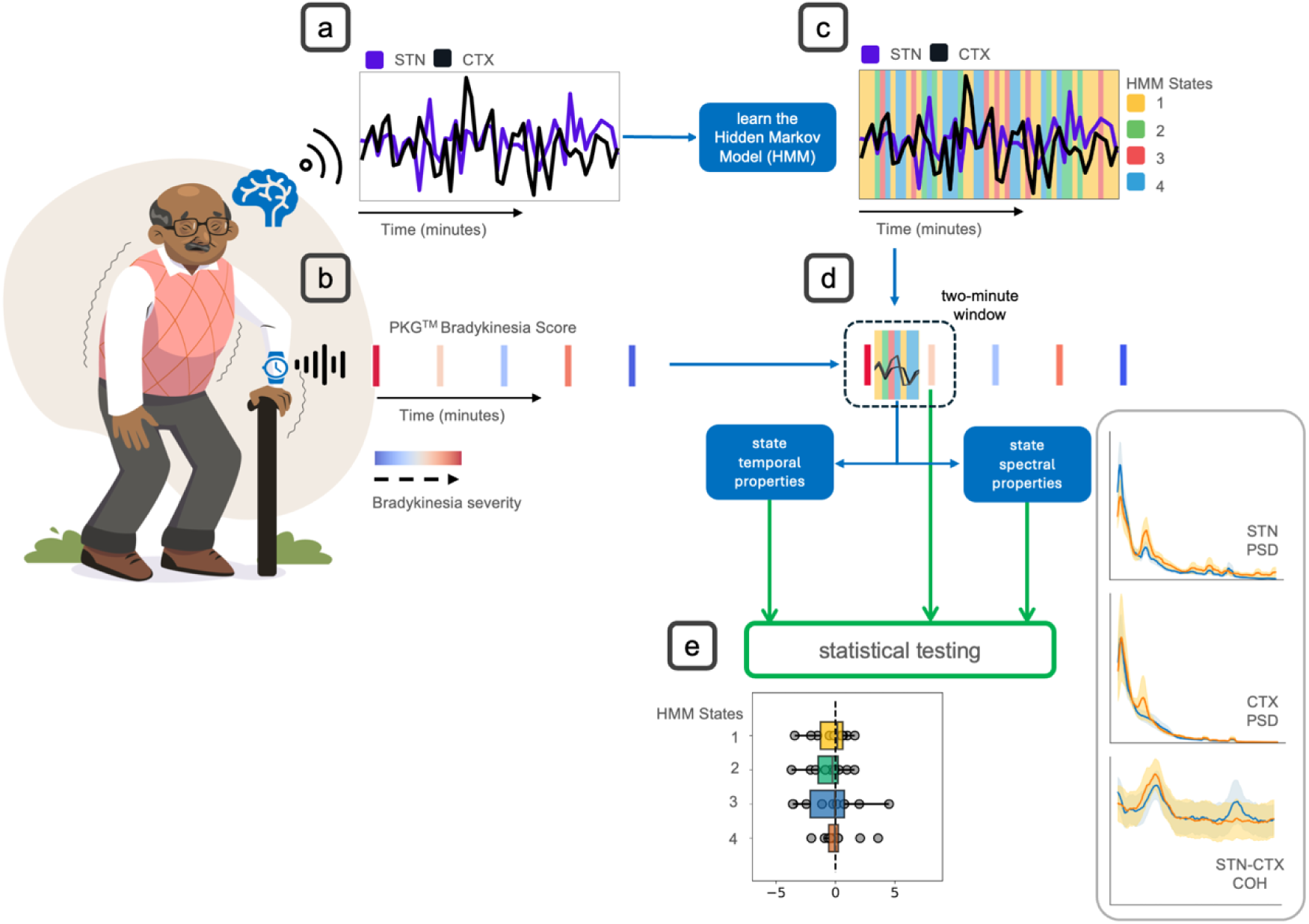
Overview of the analysis pipeline. **(a)** Neural recordings were wirelessly streamed from the subthalamic nucleus (STN) and the ipsilateral motor cortex (CTX) in Parkinson’s disease patients during naturalistic behavior. **(b)** Concurrent behavioral data were obtained via bradykinesia scores recorded every two minutes using the PKG™ watch contralateral to the hemisphere from which the neural data was streamed. **(c)** A single Hidden Markov Model (HMM) was fitted across the entire dataset, revealing distinct state time courses, representing the probability of occupying specific, statistically defined neural states at each time point. **(d)** For each two-minute window between consecutive bradykinesia scores, spectral and temporal properties were extracted for each HMM state, including power spectral density (PSD) of the STN and CTX, as well as coherence between STN and CTX. **(e)** The extracted neural features were then regressed onto the subsequent bradykinesia score, enabling an investigation of the mechanistic relationship between dynamic brain state properties and motor impairment with spatial, spectral, and temporal precision.

**Figure 2.**
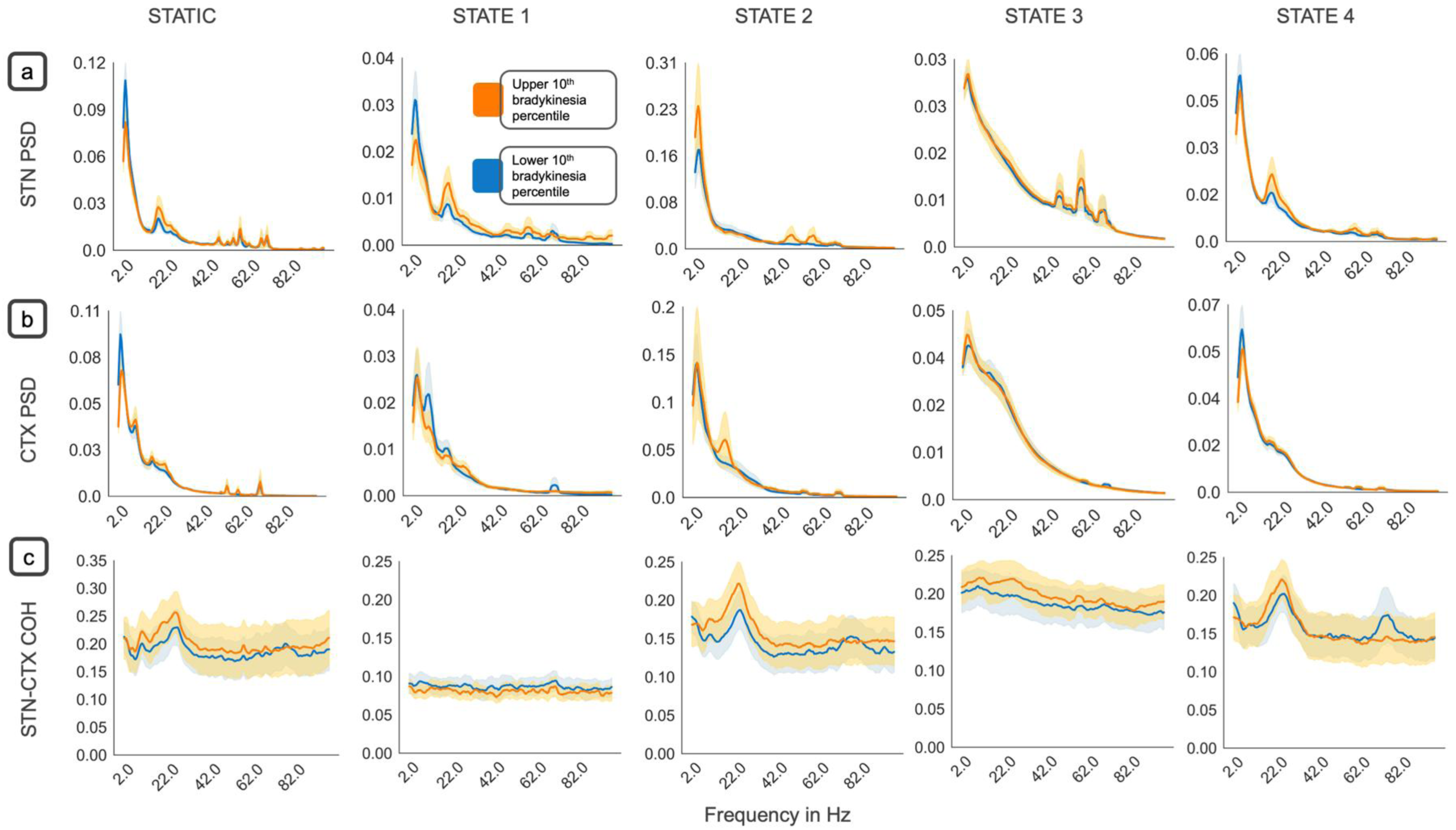
Distinct spectral fingerprints of HMM states. To investigate whether each HMM state exhibited unique or redundant spectral responses to bradykinesia, we visualised how spectral properties varied across states at different bradykinesia levels. The power spectral density (PSD) is shown for each state in **(a)** the subthalamic nucleus (STN), **(b)** the motor cortex (CTX), and **(c)** the coherence between STN and CTX. The first column displays static spectral characteristics computed across the entire dataset without state decomposition, enabling direct comparison between conventional and state-resolved spectral analysis. Spectral properties are compared between the upper 10th percentile and lower 10th percentile of bradykinesia scores. The solid line represents the mean spectra across the 5 participants and 2 hemispheres for each participant, and the shading represents the standard error of the mean across participants and hemispheres. Each state demonstrated a distinct spectral profile in response to bradykinesia. For example, **State 2** exhibited low-gamma power changes in the STN, as well as localized cortical and STN-CTX coherence alterations in the beta range with increasing bradykinesia. In contrast, **State 4** was characterized by local beta power changes in the STN, with only high-gamma coherence scaling inversely with bradykinesia at the circuit level. States 1 and 3 showed relatively flat spectral characteristics in the circuit, with no observable spectral shifts in response to bradykinesia. These findings highlight the heterogeneous neural substrates underlying bradykinesia, where some states (e.g., States 2 and 4) capture both local power and circuit-level spectral modulations, whereas others remain spectrally unresponsive to motor impairment.

To further elucidate the behavioral relevance of these spectral features, we visualized how each state’s spectral properties varied with contralateral hemibody bradykinesia severity. Given that state-wise spectra were computed over two-minute epochs, each aligned to a bradykinesia score (Methods: PKG® Alignment and Preprocessing of Behavioral Data and Spectral Characterization of HMM States), we stratified the data by bradykinesia severity. Specifically, we averaged spectral features across participants and hemispheres within the top and bottom deciles of the bradykinesia score distribution and visualized both the mean (solid line) and standard error (shaded area) for each condition. The goal of this analysis was to provide a visual intuition demonstrating that spectral characteristics differ between HMM states and systematically vary with bradykinesia severity.

The spectral architecture of each HMM state revealed distinct local and circuit-level characteristics. State 2 stood out with the highest overall power in both STN and cortical regions, yet this elevated local activity did not translate to enhanced coherence. Instead, states 2-4 all showed comparably strong STN-motor cortex coherence, while state 1 displayed markedly weaker inter-regional coupling.

Beta-band dynamics within the STN showed state-specific patterns. States 1 and 4 both exhibited clear beta peaks that grew more prominent with increasing bradykinesia severity. However, their cortical counterparts behaved differently: state 1 showed declining low-beta activity in motor cortex as bradykinesia worsened, while beta activity in state 4 remained unchanged. State 2 presented a different picture, with a cortical beta peak emerging only during high-bradykinesia periods. This divergence suggests that STN and cortical beta rhythms operate somewhat independently, with symptom sensitivity varying by network state.

Gamma frequency patterns added another layer of complexity. State 3 was distinguished by multiple gamma peaks in the STN that appeared stable regardless of bradykinesia fluctuations. States 1 and 2 also featured high-frequency STN activity, though cortical gamma remained generally subdued across most states. State 1 proved an exception, showing enhanced cortical gamma specifically during periods of reduced bradykinesia.

The coherence spectra unveiled additional state-specific circuit signatures. States 1 and 3 maintained stable broadband coherence that remained consistent across both frequencies and bradykinesia levels. In contrast, states 2 and 4 displayed more dynamic circuit-level behavior. State 2 featured a prominent high-beta coherence peak that intensified with bradykinesia severity, while state 4 showed a similar beta coherence peak that scaled less with worsening bradykinesia. Interestingly, state 4 exhibited a high-gamma coherence peak which appeared during low-bradykinesia periods, potentially representing a circuit signature of enhanced motor performance.

Taken together, these results delineate four neurophysiological states with distinct spectral and circuit-level fingerprints that scale differentially with bradykinetic severity. State 1 is characterized by reduced STN-cortical coherence and dominant STN beta activity alongside suppressed cortical beta, with this pattern intensifying during high bradykinesia periods. This suggests a disengaged state where local STN activity operates with diminished cortical synchronization. State 2 in contrast exhibits globally elevated power across regions, including increases in STN gamma and cortical beta, coupled with enhanced beta coherence. This configuration potentially marks a hyper-synchronous but maladaptive regime that coincides with poor motor function. State 3 features prominent STN gamma oscillations that remain remarkably stable across bradykinesia levels, accompanied by limited cortico-subthalamic interaction. This state appears to represent a gamma-dominant configuration largely independent of symptom fluctuations. Finally, state 4 maintains beta activity both locally in the STN and at the circuit level but uniquely expresses a high-gamma coherence peak during low-bradykinesia episodes.

This pattern highlights a potential circuit dynamic that supports improved motor control while preserving ongoing beta processes in a contextually adaptive manner. These insights provide the foundation for formal statistical testing of the relationship between bradykinesia severity and state-resolved spectral fingerprints in the following section.

### HMM States Reveal the Dynamics of Spectral Contributions to Bradykinesia

Building upon the characterization of state-specific spectral profiles, we implemented a GLM analysis to separately compare whether spectral features derived from static and HMM based analysis related to bradykinesia severity. For the static spectral analysis, contralateral bradykinesia severity served as the predictor, whilst independent variables comprised spectral power across five canonical frequency bands (delta-alpha (2-10 Hz), low-beta (12-20 Hz), high-beta (20-35 Hz), low-gamma (40-70 Hz), and high-gamma (70-100 Hz)) in three spatial domains: subthalamic nucleus (STN), motor cortex (CTX), and STN-CTX coherence.

The static spectra model explained 17% of the variance in bradykinesia scores (see left column of **Figure 3**). STN gamma activity emerged as a consistent predictor of motor impairment, with both low-gamma (t(15478) = 3.46, p < 0.01) and high-gamma (t(15478) = 3.98, p < 0.001) band activity significantly predicting increased bradykinesia. Motor cortex demonstrated an opposite pattern, with both low-gamma (t(15478) = -4.37, p < 0.001) and high-gamma (t(15478) = -4.89, p < 0.001) activity significantly associated with bradykinesia improvement. At the circuit level, delta-alpha coherence showed a significant relationship with bradykinesia improvement (t(15478) = -7.15, p < 0.001), while low-beta coherence was a significant predictor of symptom worsening (t(15478) = 8.64, p < 0.001). Interestingly neither low nor high beta power at the cortical and STN sites emerged as a significant predictor of bradykinesia severity (STN low beta, t(15478) = -0.82, p = 0.52; STN high beta, t(15478) = 2.04, p = 0.07; cortical low beta, t(15478) = 1.02, p = 0.42; cortical high beta, t(15478) = -0.66, p = 0.59).

**Figure 3.**
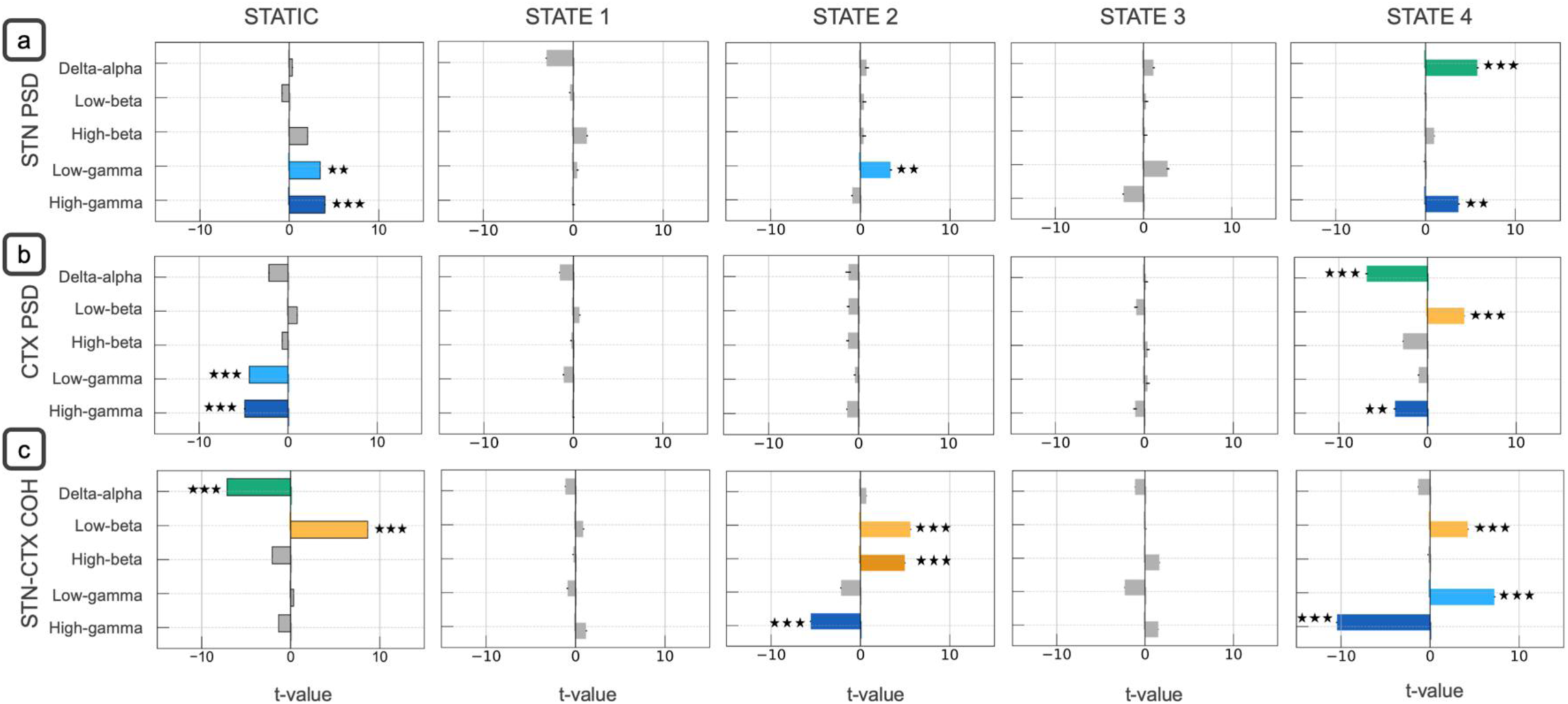
Mechanistic inference of HMM states in relation to bradykinesia severity. We applied a generalized linear model (GLM) to test how the spectral properties of each HMM-derived state predicted bradykinesia, as measured by PKG® scores. Bradykinesia severity served as the dependent variable, while predictor variables included power and coherence features from each state, computed over five frequency bands—delta–alpha (2–10 Hz), low-beta (12–20 Hz), high-beta (20–35 Hz), low-gamma (40–70 Hz), and high-gamma (70–100 Hz) at the **(a)** STN (power), **(b)** cortex (power), and **(c)** STN–cortex (coherence) levels. Hemisphere and participant identity were included as covariates. There were two hemispheres for each participant. A parallel GLM analysis using conventional static spectral features served as a baseline comparison (displayed in the first column), demonstrating that HMM decomposition revealed mechanistic insights obscured by temporal averaging. Only States 2 and 4 exhibited predictive spectral features. In **state 2**, STN low-gamma power and beta-band coherence predicted worsening bradykinesia severity. In **state 4**, STN delta–alpha and high-gamma power tracked increasing bradykinesia. In motor cortical regions for **state 4**, elevated low-beta power corresponded to worsening bradykinesia, whilst delta–alpha and high-gamma power aligned with bradykinesia ameliorations. At the circuit level, beta coherence again associated with greater impairment, while high-gamma coherence reflected reduced bradykinesia for both **state 2** and **state 4**. The horizontal box plots depict the magnitude of the t-values associated with the predictors. Each color corresponds to a different frequency band. The black error bars indicate the standard error of mean (SEM) of the t-values. Statistical significance is denoted by (***) p<0.001 and (**) p<0.01 (adjusted for multiple comparisons using Benjamini Hochberg FDR).

To establish whether HMM states reveal more nuanced spectral contributions, we implemented an analogous GLM using the same frequency bands and spatial domains, decomposed for each of the four HMM states (results presented in **Figure 3**). This state-resolved model explained 19% of the variance in bradykinesia scores. More importantly however, only two of the four states - states 2 and 4 - exhibited statistically significant associations between their spatio-spectral features and bradykinesia severity. Notably, the contributions of states 2 and 4 were mechanistically distinct and were not merely reflective of global spectral power increases.

STN low-gamma activity in state 2 was associated with worsening motor symptoms (t(15433) = 3.42, p < 0.01). In state 4, high-gamma (t(15433) = 3.71, p < 0.01) and delta-alpha activity (t(15433) = 5.81, p < 0.001) in the STN similarly predicted increased bradykinesia severity. At the cortical level, only state 4 exhibited significant associations between local spectral activity and bradykinesia severity. Increased cortical power in the delta-alpha (t(15433) = -6.89, p < 0.001) and high-gamma bands (t(15433) = -3.68, p < 0.01) correlated with motoric improvement, whereas elevated low-beta power was associated with symptom exacerbation (t(15433) = 4.20, p < 0.001).

Circuit-level interactions revealed that beta-band coherence consistently correlated with increased bradykinesia severity in both states 2 (low-beta: t(15433) = 5.63, p < 0.001; high-beta: t(15433) = 5.01, p < 0.001) and state 4 (low-beta: t(15433) = 4.26, p < 0.001). In state 2, this pathological coherence emerged without corresponding local beta associations with bradykinesia worsening in either cortex or STN. In contrast, state 4’s beta coherence was accompanied by elevated cortical low-beta activity that was also linked to motor impairment. High-gamma coherence in both states 2 (t(15433) = -5.60, p < 0.001) and 4 (t(15433) = -10.48, p < 0.001) was associated with reduced bradykinesia, although low-gamma coherence in state 4 predicted worsening symptoms (t(15433) = 7.25, p < 0.001).

The transition from conventional spectral analysis to state-resolved analysis revealed several mechanistic insights that were obscured by temporal averaging. In the static analysis, both STN low-gamma and high-gamma activity predicted worsening bradykinesia. The HMM decomposition revealed that these gamma components do not act together within the same state. When high-gamma activity strongly predicted bradykinesia locally in the STN, it emerged alongside pathological delta-alpha oscillations, which were absent in the static analysis. Cortical low-gamma activity emerged as a protective factor in the static analysis but did not emerge as a significant predictor in any individual state. High-gamma activity emerged as a significant protective factor only in state 4, where low-frequency delta-alpha activity also emerged, mirroring STN activity but with opposite clinical effects.

Delta-alpha coherence emerged as a significant protective factor in the static analysis but did not emerge as a significant factor in any individual state, suggesting that its protective effect was distributed across local cortical and STN components rather than circuit-level synchronization. Importantly, low-beta circuit coherence, which predicted worsening bradykinesia in the static analysis, emerged only in two states while maintaining its association with symptom deterioration. Additional circuit markers emerged exclusively in the state-resolved analysis, including high-gamma coherence in both states 2 and 4 (predicting bradykinesia improvement), low-gamma coherence as a marker of worsening bradykinesia in state 4, and high-beta coherence in state 2 also predicting symptom worsening.

Taken together, these findings emphasize the state dependence of the neural correlates of bradykinesia, which encompass both spectrally and spatially dissociable mechanisms. Importantly, spectral changes within a specific anatomical region do not necessarily coincide with analogous changes at the circuit level. While STN-based spectral features were uniformly associated with worsening motor symptoms, the motor cortex and STN-CTX circuit exhibited more nuanced and state-dependent spectral architectures, offering potentially richer biomarkers for tracking and therapeutic intervention.

### Temporal Dynamics Reveal a Compensatory Beta State

A key advantage of HMMs is their capacity to extract both temporal and spatio-spectral properties of latent neural states. We computed temporal metrics for each state (fractional occupancy, mean state lifetime and inter-visit interval) in order to establish their relationship to bradykinesia severity (see Methods: Temporal Metrics of HMM States).

Interestingly, states 2 and 4, which previously exhibited distinct coherence signatures in the spectral domain, demonstrated the highest fractional occupancies, indicating that these two network configurations dominated the neural time series relative to states 1 and 3 (**Figure 4a**). In parallel, these same states (2 and 4) also displayed prolonged average lifetimes (**Figure 4b**). Notably, only state 3 exhibited extended inter-visit intervals, implying more sporadic activation (**Figure 4c**). Individual participants displayed idiosyncratic trends in temporal features ordered by the upper and lower bradykinesia deciles (as illustrated in the paired line plots, **Figures 4a-d**).

**Figure 4.**
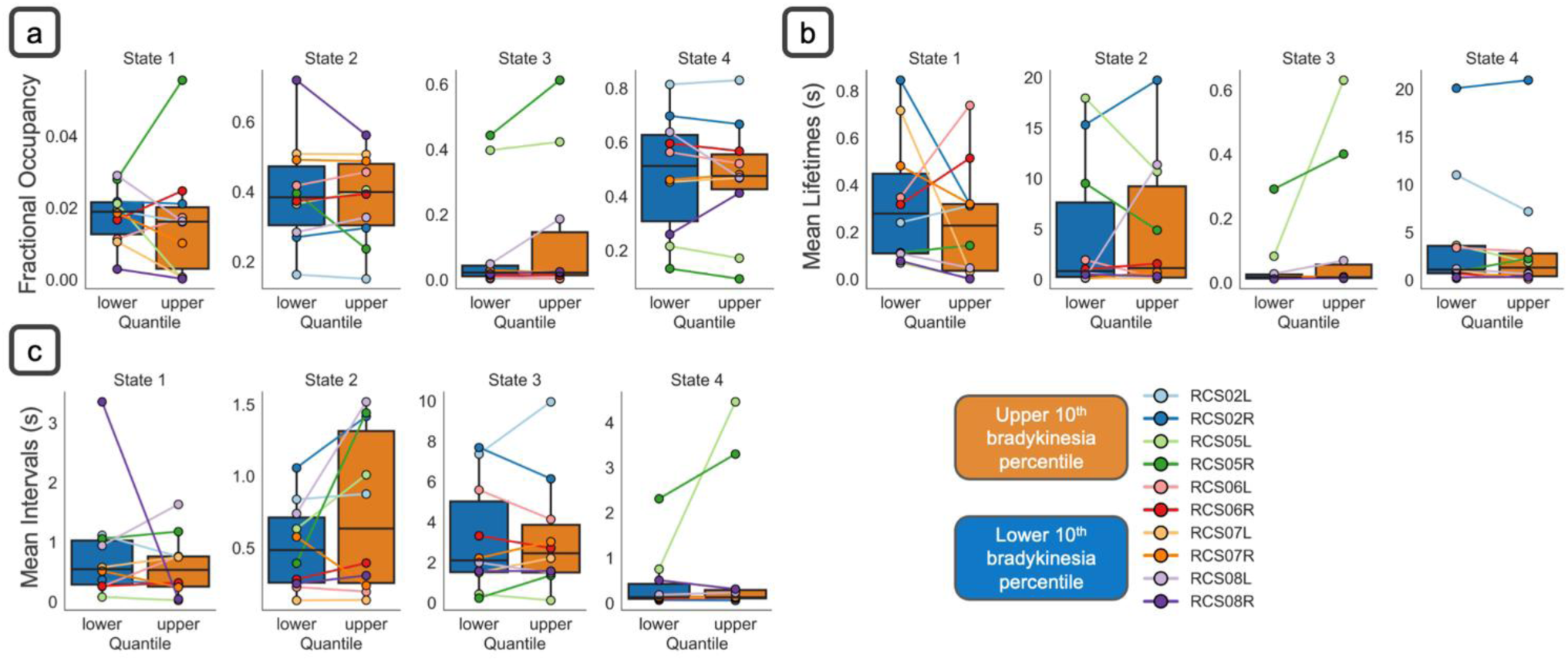
Temporal dynamics of HMM states in relation to bradykinesia. In addition to their spectral properties (Figure 2), HMM states exhibit distinct temporal characteristics that vary with bradykinesia severity. We quantified three key temporal metrics: (a) **Fractional Occupancy (FO):** The proportion of time each state is active within the dataset. (b) **Mean Lifetime:** The average duration a state remains continuously active before transitioning. (c) **Mean Interval Visit:** The time interval between consecutive activations of the same state. Each metric was computed for all four HMM states, comparing data from the upper and lower 10th percentiles of the bradykinesia score. In the box plots, the line represents the mean and error bars indicate the standard error of the mean (SEM). Individual participant-hemisphere combinations are also shown as connected line plots across the two bradykinesia levels, with colors corresponding to individual participants as detailed in the figure legend. Our findings reveal that the two states exhibiting spectrally distinct circuit-level responses to bradykinesia (**States 2** and **4**) were the most dominant across participants. These states were active for longer durations, persisting from several seconds to tens of seconds. In contrast, the two states that lacked circuit-level spectral responses to bradykinesia (**States 1** and **3**) were short-lived (hundreds of milliseconds) and less prevalent across participants. These results highlight the differential engagement of dynamic neural states in the pathophysiology of bradykinesia.

GLM analysis (see Methods: Modelling Bradykinesia via Temporal Features) revealed that state 1 exhibited temporal properties consistently associated with improved motor function. Specifically, increased fractional occupancy (t(15492)=-3.81;p-value<0.001) and prolonged lifetimes of state 1 (t(15489)=-3.65;p-value<0.001) were significantly associated with reduced bradykinesia severity, suggesting that greater temporal persistence of this state predicts motor improvement (**Figure 5a**, **5c**). Longer inter-visit intervals for state 1 were also linked to bradykinesia relief (t(15489)=-3.36;p-value<0.001), indicating that temporally dispersed, but sustained re-engagements with this state may be beneficial (**Figure 5b**). Importantly, although state 1’s spectral features were not directly predictive of bradykinesia, it did exhibit peaks of beta activity both within the cortex and within the STN (**Figure 2a**, **2b**; state 1) that were modulated by symptom severity and not associated with cortico-STN beta coherence peaks (**Figure 2c**).

**Figure 5.**
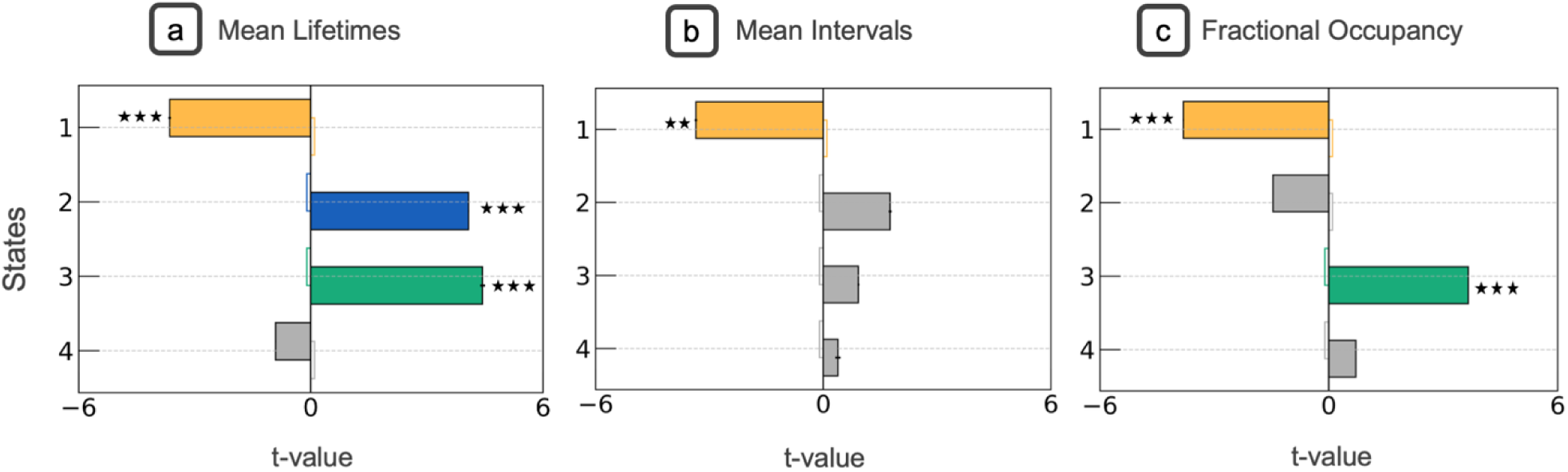
Temporal metrics of HMM-derived neural states as predictors of bradykinesia severity. We employed a general linear model (GLM) framework to assess how the temporal characteristics of each latent neural state predicted bradykinesia severity, as indexed by PKG® scores. Bradykinesia served as the dependent variable, and independent variables included state-specific temporal features: fractional occupancy, mean lifetime and mean inter-visit interval. Each temporal metric was evaluated in a separate GLM to isolate its individual contribution. See Methods for detailed modeling procedures. Increased fractional occupancy and prolonged lifetimes of **state 1** were significantly associated with reductions of bradykinesia, indicating that greater temporal persistence of this state predicts improved motor function. Interestingly, longer inter-visit intervals for **state 1** also predicted bradykinesia relief, suggesting that temporally dispersed but sustained re-entries into this state were beneficial. Conversely, temporal features of **states 2** and **4** were associated with greater symptom severity. Together, these findings highlight that not only the presence but also the temporal architecture of specific brain states contributes meaningfully to bradykinesia dynamics. The horizontal box plots depict the magnitude and signage of the t-values associated with the predictors. Each color corresponds to a different state. The black error bars at the end of the bar plots depict the standard error of mean (SEM) of the t-values. Statistical significance is denoted by (***) p<0.001 and (**) p<0.01 (adjusted for multiple comparisons using Benjamini Hochberg FDR).

Our analysis also revealed that increased fractional occupancy (t(15492)= 3.66;p-value<0.001) and prolonged lifetimes (t(15489)= 4.44;p-value<0.001) of state 3 were both associated with greater bradykinesia severity (**Figure 5a**, **5c**). Interestingly, this effect occurred in the absence of a correlation between state 3’s spectral features and bradykinesia, and despite there being no observable scaling of this state’s spectral profile with bradykinesia severity across deciles. Finally, we also observed that increased lifetimes of state 2 – a state that predominantly exhibited spectral features correlating with worsening bradykinesia - were associated within increased bradykinesia severity (t(15489)= 4.07;p-value<0.001).

Collectively we conclude that temporal dynamics of latent brain states are important predictors of motor impairment. While states 2 and 4 are temporally dominant, it is predominantly their internal spectral features that associate with bradykinesia severity. Surprisingly, state 1, which is characterised by short-lived STN and motor cortical beta activity, may represent a compensatory motoric configuration that ameliorates bradykinesia. This suggests that relief from bradykinesia may depend not only on suppressing pathological states, but also on increasing the occurrence of compensatory configurations.

## Discussion

We leveraged Hidden Markov Models to discover latent neural states of cortical and STN activity in patients with Parkinson’s disease (PD). By relating these identified neural states to dynamic measurements of bradykinesia, we show that bradykinesia has a multidimensional nature, encoded through both the spectral and temporal properties of states of cortico-STN network activity. Our analysis speaks to the utility of moving beyond single site and single frequency band biomarkers (e.g., STN beta activity), towards a more nuanced understanding of how moment-to-moment fluctuations in brain network dynamics underpin PD pathophysiology.

A key finding of this work is the existence of a compensatory state characterised by short-lived, but uncoupled STN and cortical beta activity that associates with ameliorations of bradykinesia. This compensatory state could represent a potential target state for neuromodulation therapies, and its presence is supported by previous studies demonstrating that the occurrence of short-lived bursts of beta activity within the STN can be associated with improved motor performance^49^. Moreover, adaptive DBS approaches which act over slower timescales to potentially preserve physiological levels of beta within motor circuits could be more effective than approaches which seek to rapidly terminate beta activity^50^.

In what follows, we further discuss our findings with reference to the existing literature regarding the roles of beta and gamma band activities in PD pathophysiology.

### Beta Oscillations: State Dependent Roles in Bradykinesia

Beta oscillations (13–30 Hz) have been a central focus in PD research, due in large part to their consistent elevation in the subthalamic nucleus (STN) and cortex under dopamine-depleted conditions^7–9^. Increased STN beta power has been linked to bradykinesia and rigidity, prompting clinical strategies such as beta-guided adaptive DBS, that aim to reduce beta amplitude^51^. Our findings nuance this established paradigm by demonstrating that STN beta power alone was not an independent predictor of bradykinesia severity. Instead, beta-related pathology emerged most strongly through circuit-level features, especially beta-band coherence between the STN and motor cortex. This distinction underscores the importance of network synchrony over local oscillatory magnitude in driving motor impairment.

A more granular analysis of the state-specific data revealed that cortical beta activity had a greater predictive influence on motor impairment than subthalamic nucleus (STN) beta power in certain HMM states. For example, in state 4, low-beta activity in the motor cortex emerged as a robust predictor of bradykinesia severity, whereas in other states (e.g., state 2), neither STN nor cortical beta power significantly contributed to bradykinesia.

State 1 offered additional insights, particularly with respect to temporal dynamics. State 1 exhibited a distinct beta peak within the STN across both low and high bradykinesia deciles, with a slightly lower frequency beta peak localized to the motor cortex only in the low bradykinesia decile. Spectrally, state 1 exhibited a relatively flat profile of circuit level coherence. Although the spectral features of state 1 were not associated with bradykinesia, increased temporal occurrence of state 1 was predictive of bradykinesia ameliorations.

Taken together, these findings underscore the state dependent role of beta oscillations, wherein they are not inherently detrimental, but become pathologically relevant when occurring within certain temporally segregated network configurations.

### Gamma Oscillations: State Dependent and Spatially Specific effects on Bradykinesia

A growing literature frames gamma-band activity (50–100 Hz) as broadly “prokinetic” in PD, particularly because it is enhanced by dopaminergic medications and correlates with improved movement speed ^39–42^. Our results generally support this prokinetic narrative for STN–cortical high gamma coherence (70-100 Hz), yet they also highlight a crucial caveat: local gamma oscillations within the STN can, in some states, associate with worsening bradykinesia. This contrast underscores the multifaceted role of gamma band oscillations, whereby their functional impact depends on: (1) the site where gamma emerges (cortex vs. STN), (2) whether it is expressed coherently at the circuit level or remains an isolated local burst, and (3) the nature of lower frequency oscillations that emerge as significant predictors in the same spatial location.

### The Paradox of Local STN Gamma vs Motor Cortex Gamma

In state 4, local STN high-gamma power scaled positively with bradykinesia severity. One particularly telling pattern was the shared signage of high-gamma and low-frequency (delta– alpha, ∼2–10 Hz) features in our regression models. That is, when STN high-gamma increased in these states, STN delta-alpha power rose in parallel, and both predicted greater motoric impairment (**Figure 3a**; state 4). Interestingly, our data revealed contrasting results for the motor cortex in state 4, where increased local cortical high-gamma and delta-alpha activity were associated with reduced bradykinesia scores (**Figure 3b**; state 4).

The tight relationship between low frequency delta/alpha oscillations and gamma, and their shared differential impact on bradykinesia at the cortical and STN sites, raises the possibility of local non-linear mechanisms for propagating gamma oscillations^28,52–56^. For example, high gamma in the STN could be phase locked to or entrained by slower local STN rhythms often associated with akinetic or low dopaminergic states^33^. Such coupling could neutralize or reverse gamma’s usual prokinetic advantage by aligning it with activity that can perpetuate bradykinesia.

### Circuit-Level Gamma Coherence as a Prokinetic Marker

STN–cortical high gamma coherence exerted prokinetic influences across states in our data. This distinction highlights that gamma coherence, rather than local gamma amplitude alone may be the key signal for binding cortex and STN into a functional network that can initiate movements efficiently. Prior work^40,41^ has shown that dopaminergic medication and deep brain stimulation enhance cortico-subthalamic gamma coupling at movement onset, particularly during faster or larger-amplitude movements. Our findings bolster that narrative but add nuance by showing how coherence can remain prokinetic (**Figure 3b**; state 2) even when local STN gamma amplitude might be anti-kinetic (**Figure 3a**; state 2). Functionally, this suggests that the network requires high-frequency communication channels to bypass or counteract pathological beta synchrony; in the absence of that coherent linkage, local STN gamma by itself does not guarantee movement facilitation.

### Clinical implications and Study Limitations

Taken together, our observations underscore that beta amplitude alone, particularly in the STN, is insufficient as a biomarker of bradykinesia. Beta-band coherence may offer a more reliable gauge of pathological state transitions. Consequently, therapies that aim solely to suppress beta power may yield inconsistent outcomes if they ignore the underlying circuit configuration and neglect compensatory beta states that associate with bradykinesia ameliorations. State-aware neuromodulation could specifically target neural states in which beta coherence is most detrimental, potentially allowing for a more precise and adaptive approach to bradykinesia relief.

Similarly for gamma oscillations, interventions that non-specifically “enhance gamma” (e.g., via transcranial stimulation) could yield inconsistent results if they fail to target coherent gamma bursts or if they inadvertently exacerbate local STN gamma tied to pathological states. Looking ahead, adaptive DBS systems might deliver stimulation only when gamma coherence dips or when local STN gamma uncouples from cortical input. Such closed-loop methods could selectively reinforce the circuit-level gamma that has prokinetic effects. Looking ahead, future neuromodulation strategies might dynamically track multiple spectral and temporal features (rather than a single “beta amplitude” threshold) to disrupt harmful synchrony while sparing or fostering beneficial oscillatory states.

Several study limitations merit consideration. Firstly, we utilized a four-state HMM, selected for its balance of interpretability and explanatory power. However, this may underrepresent the full spectrum of dynamic states present in the data. Future models could explore larger or hierarchical state architectures or incorporate explicit duration modelling to capture the persistence of pathological versus compensatory states. Secondly, while PKG® sensors provided continuous, ecologically valid measures of motor function, their resolution is limited compared to high-density motion capture or wearable kinematic systems. This constraint may obscure finer resolution motor fluctuations and their alignment with neural states. Finally, the correlational nature of our analysis limits causal interpretation. Real-time, state-triggered neuromodulation experiments are necessary to determine whether selectively modulating specific neural states can causally alter PD symptoms.

## Conclusions

Our study provides high spatial and temporal resolution insights into the link between cortico-STN network activity and bradykinesia in PD. By modelling brain activity as transitions between statistically defined states with distinct spectral and coherence profiles, we capture a dynamic view of symptom expression that enables us to move beyond time-averaged biomarkers. This state-based approach could offer practical pathways toward adaptive, precision-guided interventions that respond to how the brain organizes its own activity in real time.

## Contributors

Conceptualization, A.S., W.N., S.L., P.S., A.O.; Methodology, A.S., A.H., M.S., S.L., P.S., A.O.; Software & Formal Analysis, A.S., T.L., B.A.; Data Curation, A.S., A.H., M.S., W.N., S.L., P.S., A.O.; Supervision, S.L., P.S., A.O.; Funding Acquisition, S.L., P.S., A.O.

## Declaration of Interests

P.S. is a consultant for Neuralink and InBrain Neuroelectronics. S.L. is a consultant for Iota Biosciences. The other authors declare no competing interests.

## Data sharing statement

Anonymised patient data are available from the corresponding author on request.

---

**Supplementary Figure 1.**
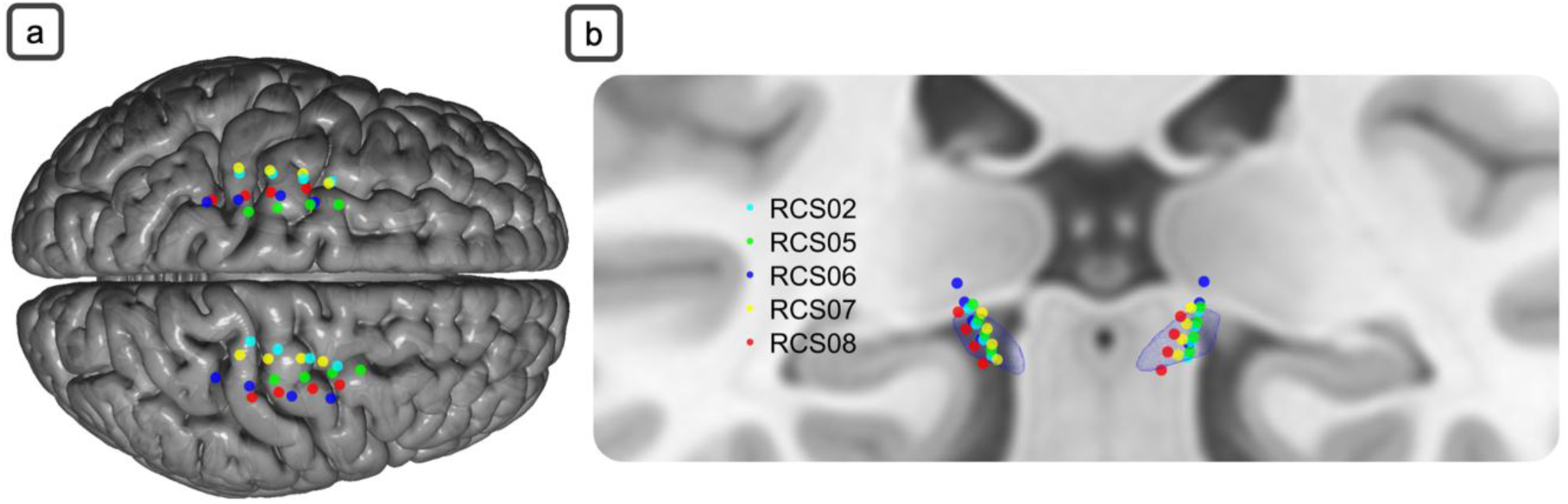
Electrode contact locations. **(a)** The left image shows cortical contact locations, superimposed on a cortical mesh derived from the same template MRI. (b) The right image shows STN electrode contact locations for all 5 patients, superimposed on an STN mesh (blue) and template MRI (coronal view) in Montreal Neurological Institute (MNI) space.

